# Telomere, epigenetic clock, and biomarker-composite quantifications of biological aging: Do they measure the same thing?

**DOI:** 10.1101/071373

**Authors:** D.W. Belsky, T.E. Moffitt, A.A. Cohen, D. Corcoran, M.E. Levine, J. Prinz, J. Schaefer, K. Sugden, B. Williams, R. Poulton, A. Caspi

## Abstract

The geroscience hypothesis posits that therapies to retard biological processes of aging can prevent disease. To test such “geroprotective” therapies in humans, surrogate endpoints are needed for extension of disease-free lifespan. Methods to quantify biological aging could provide such surrogate endpoints, but different methods have not been systematically evaluated in the same humans. We studied seven measures of biological aging in 964 middle-aged humans in the Dunedin Study: telomere-length, three epigenetic-clocks, and three biomarker-composites. Agreement between these different measures of biological aging was low. We also compared associations between biological aging measures and outcomes that geroprotective therapies will seek to modify: physical functioning, cognitive decline, and subjective signs of aging. The 71-CpG epigenetic clock and the biomarker composites were consistently related to these outcomes. Effect-sizes were modest. Quantification of biological aging is a young field. Next steps are to move toward systematic evaluation and refinement of methods.

## INTRODUCTION

Data syntheses in biodemography and gerontology identify aging as the leading cause of human morbidity and mortality^1,2^. The so-called “geroscience hypothesis” builds on these data to posit that interventions to slow the biological processes of aging could prevent or delay many different diseases simultaneously, prolonging the healthy years of life^3^. Econometric projections suggest interventions that achieve even modest slowing of biological aging could reduce burden of disease more than curing all cancer and heart disease combined^4^. Candidate interventions to slow aging are emerging from studies of animals^5,6^. However, a barrier to translating these models to help humans is that human aging is a gradual, slow-moving process that is not easily measured in clinical trials. Observing completed human lifespans or even healthspans (the portion of lifespan preceding onset of chronic disease) is time- and cost-prohibitive. In order to refine intervention targets and evaluate intervention effectiveness, surrogate endpoints are needed that can stand in as proxies for extended lifespans or healthspans^7^. Thus, quantifications of biological aging are of growing interest in biomedical and social sciences.

Measures of biological aging are intended to provide proxy measurements of lifespan/healthspan. In contrast to chronological age, which increases at the same rate for everyone, biological aging can occur at different rates in different individuals. Various measures of biological aging have been proposed, including telomere length, algorithms applied to genome-wide DNA methylation data, and algorithms combining information on multiple clinical biomarkers^8–14^. However, it is not known if these various approaches to quantifying biological aging measure the same or different aspects of the aging process. In addition, it is unknown if some proposed methods are more closely associated with healthspan than others.

We conducted a study of seven prominent methods to quantify biological aging in a 1-year birth cohort of 1,037 adults followed prospectively to midlife with 95% retention: the Dunedin Study. When cohort members turned 38 years, approximately the midpoint of the contemporary human lifespan, we assessed their biological age using seven different methods. Although there is emerging evidence for each of these methods, there have not been studies to evaluate all of them simultaneously in the same group of humans. We tested if the different measures quantified the same aging process by computing correlations among the different biological aging measures. A second issue is that although biological age measured in later life has been shown to predict disease and mortality, it is unknown if biological age measured in midlife can predict healthpsan. To extend helathspan, “geroprotective” therapies must be delivered prior to the onset of disease and disability, i.e. in people who are still relatively young and healthy. Validation is therefore needed in this younger population to establish proof of concept that biological aging measures can serve as surrogate endpoints for healthspan extension in clinical trials of geroprotective therapies. One form of construct validation that has been proposed for evaluating new measures of biological aging is to test correlation with chronological age. We extended this construct validation to test signs of aging in a group of humans who are all the same chronological age. We analyzed signs of aging that geroprotective interventions will aim to ameliorate: worsening physical functioning, cognitive impairment and decline, and subjective perceptions of declining health.

## RESULTS

Biological aging measures can be divided along three axes. One axis is the technical dimension of the number of assays required (e.g., telomere length is measured with a single assay whereas multiple assays are required for algorithms that combine different types of biomarkers). A second axis is the measurement design (i.e., a single cross-sectional measurement versus repeated, longitudinal measurements). A third axis is the biological level at which measures are implemented (e.g. telomeres are a cellular-level measure typically implemented in a specific tissue whereas multi-biomarker algorithms are patient-level measures that combine information from multiple organ systems). We implemented seven methods to quantify biological aging using data from the Dunedin Study Biobank. These measures are grouped according to the three axes in Table 1.

**Table 1.** Taxonomy of Biological Aging Measures for Use in Humans Evaluated in this Article.

| Variety of Assays |  | Measurement Design |  | Biological Level of Measurement |
| --- | --- | --- | --- | --- |
|  |  | Cross-sectional | Longitudinal |  |
|  | **Single Assay** | Telomere Length, Epigenetic Clocks* | Telomere Erosion, Epigenetic Ticking* | Cellular-level measures assayed in blood |
|  | **Multiple Assays** | KDM Biological Age, Age-related Homeostatic Dysregulation | Pace of Aging | Patient-level measures derived from assays of multiple organ systems |

\*Epigenetic clocks are composed of dozens or hundreds of different methylation marks across the genome. We classify the clocks in the “single measure” row because genome-wide DNA methylation is measured in a single assay and reflects a single biological substrate.

Telomere length and epigenetic clocks have been proposed as cross-sectional estimates of biological aging based on a single biological measure (Table 1, top left panel).

We measured leukocyte <u>telomere length</u> from peripheral-blood DNA using the qPCR method^15^. At age 38, telomere length was approximately normally distributed in the cohort (T/S ratio M=1.05, SD=0.32)

We measured three different <u>epigenetic clocks</u> based on 353^9^, 99^13^, and 71^10^ CpG sites, respectively, from whole-genome DNA methylation assayed from peripheral-blood DNA using Illumina 450k chips (Illumina Inc., CA, USA). At age 38, epigenetic clocks were approximately normally distributed in the cohort and accurately centered on the observed chronological age (for the for the 353-CpG Clock, M=37y, SD=4; for the 99-CpG clock M=38y, SD=5; for the 71-CpG clock M=37y, SD=5).

In addition to these age-38 measurements, we also measured Study members’ telomere length and epigenetic clock values from blood samples taken when they were aged 26 years. We calculated longitudinal <u>telomere erosion</u> and <u>epigenetic ticking rates</u> by subtracting age-26 values from age 38 values (Table 1, top right panel).

Klemera-Doubal-method Biological Age^16^ and Age-related Homeostatic Dysregulation^17^ have been proposed as cross-sectional estimates of biological aging based on multiple biological measures (Table 1, bottom left panel).

We measured <u>Klemera-Doubal-method Biological Age</u> (hereafter “KDM Biological Age”) from 10 blood- and organ-system-function biomarkers assessed using standard assays. At age 38, KDM Biological Age was approximately normally distributed in the cohort (M=38y, SD=3).

We measured <u>Age-related Homeostatic Dysregulation</u> from 18 blood- and organsystem-function biomarkers assessed using standard assays. This measure quantifies deviation from a reference norm in Mahalanobis distance^18^. We used the normative values for the Dunedin cohort when they were aged 26 years to form this reference. We log transformed the computed distances for analysis. At age 38, Age-related Homeostatic Dysregulation was approximately normally distributed in the cohort (M=3.37, SD=0.61).

<u>Pace of Aging</u> is a longitudinal estimate of biological aging based on changes across repeated measurements of multiple biological measures (Table 1, bottom right panel). We measured Pace of Aging from changes in 18 blood- and organ-system-functional biomarkers assayed when Study members were aged 26, 32, and 38 years^14^. Pace of Aging quantifies the rate of biological aging in years-of-physiological-change-per-chronological-year units (M=1, SD=0.38).

In summary, by midlife, members of the Dunedin cohort, who were all born in a 1-year period and followed up 38 years later, varied on all seven estimates of biological age. Distributions of biological age estimates are shown in Figure 1.

**Figure 1.**
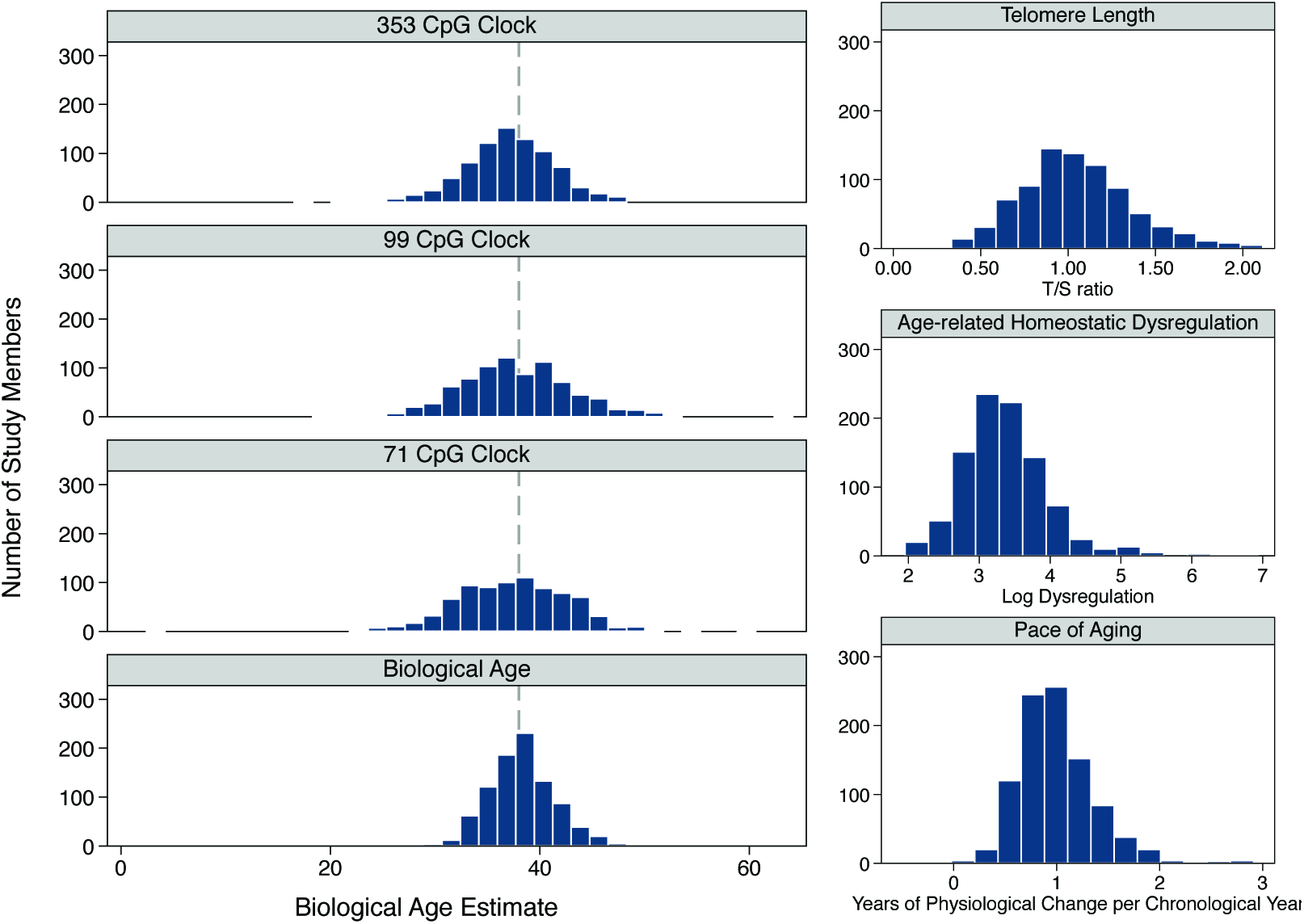
Distributions of cross-sectional biological aging measures and Pace of Aging in the Dunedin birth cohort at age 38 years. The left side of the figure plots estimated biological ages based on the 353-, 99-, and 71-CpG epigenetic clocks and the Biological Age algorithm. The dashed gray line shows age-38 years, the chronological age of the cohort at the time assays were taken. The right side of the figure plots values for telomere length Age-related Homeostatic Dysregulation, also assayed at chronological age 38 years, and Pace of Aging, which was derived based on repeated measurements taken at ages 26, 32, and 38 years.

### Do proposed methods to quantify biological aging measure the same features of the aging process?

To test the hypothesis that the different biological aging measures quantify the same aging process, we computed correlations among the different measures (Figure 2). Telomere, epigenetic clock, and clinical-biomarker algorithm methods showed little overlap. Epigenetic clocks were correlated with each other in the r=0.3-0.5 range. Clinical biomarker algorithm measures were correlated with one another in the r=0.4-0.6 range. However, telomere length was not significantly correlated with estimates from epigenetic clocks or clinical-biomarker algorithms (|r|=0.02-0.05) and correlations of epigenetic clock measures with correlations of clinical-biomarker-algorithm measures with epigenetic clock measures were generally low. The 71-CpG clock, was weakly correlated with the clinical biomarker measures (r=0.10-0.15, p<0.001 for all) and the 353- and 99-CpG clocks were also weakly correlated with KDM Biological Age (r=0.07-0.08, p<0.05 for both). Results were similar when Spearman correlations were computed to reduce the influence of extreme values (Supplemental Table 1).

**Figure 2.**
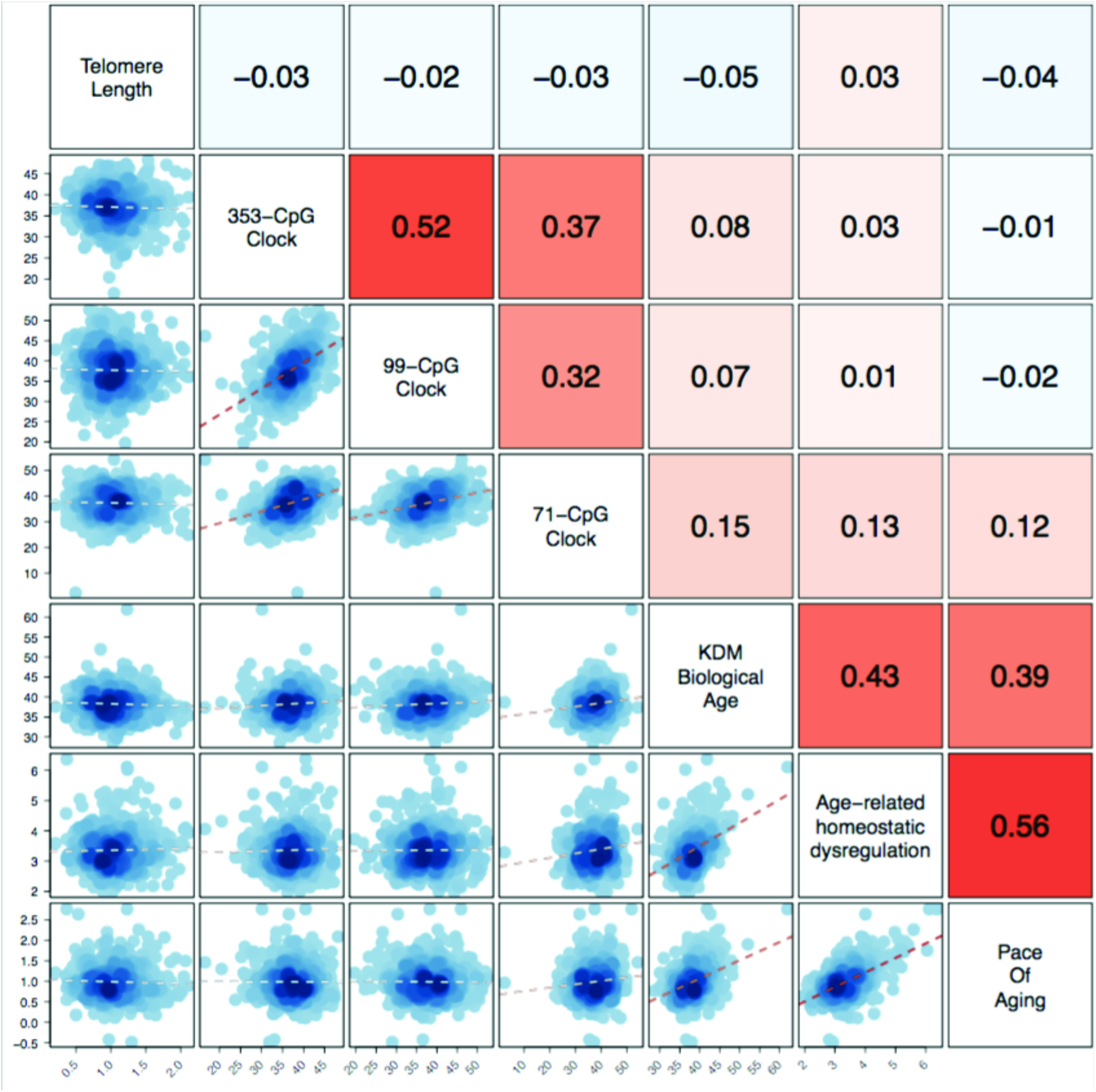
Relationships among seven measures of biological aging in a birth cohort at chronological age 38 years. The figure shows a matrix of scatterplots and correlations illustrating relationships among seven measures of biological aging: Leukocyte telomere length, 353-, 99-, and 71-CpG epigenetic clocks, KDM Biological Age, Age-related Homeostatic Dysregulation, and Pace of Aging. Data are for n=800 Study members with complete data on all biological aging measures. Correlations are shown above the diagonal. (Correlations ≥0.07 are statistically significant at p<0.05.) Scatter plots are shown below the diagonal. Y-axis scales correspond to the biological aging metric listed to the right of the plot. X-axis scales correspond to the biological aging metric listed above the plot.

### Do proposed methods to quantify biological aging predict differences in healthspan-related characteristics at midlife?

We next tested if different methods provided comparable information about healthspan. For each biological aging measure, we tested associations with three groups of healthspan-related measures: First, we tested if biological aging measures predicted deficits in physical functioning by examining Study members’ performance on tests of balance, grip strength, and motor coordination, and by interviewing Study members about any physical limitations in carrying out activities in their daily lives. Second, we tested if biological aging measures predicted earlyonset cognitive decline by comparing Study members’ scores on cognitive tests taken at midlife to scores on parallel tests that they took when they were children. Third, we tested if biological aging measures predicted subjective signs of aging, which we measured by interviewing the Study members themselves and from observer-ratings of the Study members’ aged appearance based on facial photographs.

Telomere length was not statistically associated with healthspan-related characteristics, with the exception of facial aging (r=0.07). Likewise, the 353- and 99-CpG clocks were not statistically associated with healthspan-related characteristics (p>0.05 for all). However, older epigenetic age measured by the 71-CpG clock was statistically associated with poorer healthspan-related outcomes in all cases except for grip strength (0.05≤|r|≤0.16). The three clinical biomarker algorithms were all statistically associated with poorer healthspan-related characteristics (0.10≤|r|≤0.20 for most analyses), with the exception that Age-related Homeostatic Dysregulation was not statistically associated with grip strength. Effect sizes for healthspan-related characteristics are graphed in Figure 3. Effect sizes for subtests of cognitive function and cognitive decline are graphed in Supplemental Figure 1.

**Figure 3.**
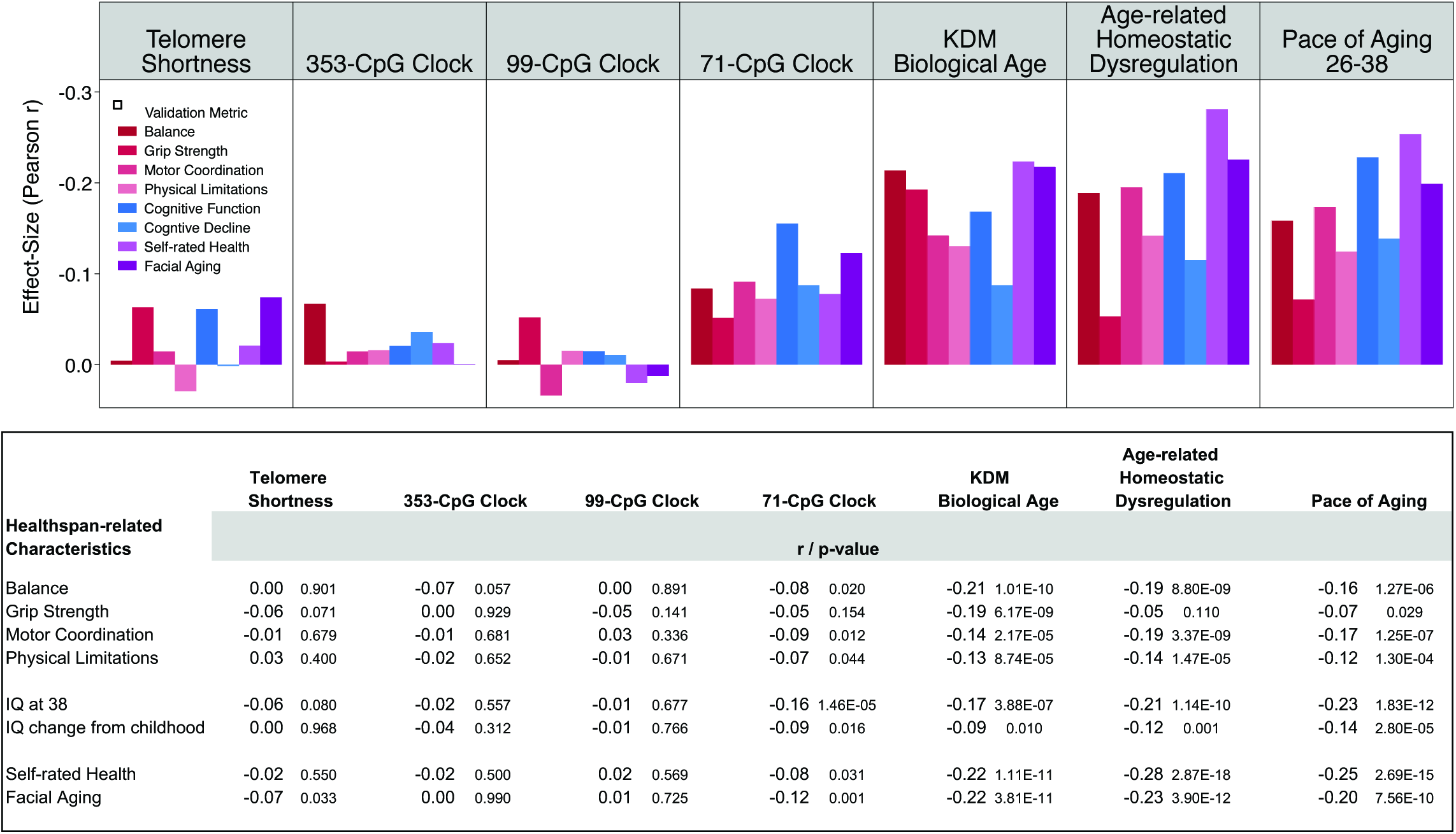
Associations of cross-sectional biological aging measures and Pace of Aging with healthspan-related characteristics. The figure shows bar charts of effect-sizes for each of the seven measures of biological aging. Effect-sizes were estimated for four measures of physical functioning (balance, grip strength, motor coordination, and self-reported physical limitations), cognitive functioning (IQ score at age 38 from the Wechsler Adult Intelligence Scale), cognitive decline (change in Wechsler-scale IQ score since childhood), and two measures of subjective aging (self-rated health and facial aging from assessments of facial photographs of the study member by independent raters). Validation metrics were scored so that higher values indicated increased healthspan. Telomere length was reversed for this analysis so that higher values corresponded to shorter telomeres. Thus, the expected direction of association for all correlations was negative—because faster biological aging is expected to shorten healthspan. Standardized regression coefficients (interpretable as Pearson r) and their p-values are reported in the table below the figure.

### Does change between repeated cross-sectional measures of biological aging track the aging process?

Most methods to quantify biological aging are designed for implementation in a cross-section of biomarker data. These cross-sectional methods could be used to measure changes in the rate of aging caused by geroprotective intervention if they were repeated, for example before and after administration of therapy. We were able to test if cross-sectional biological-age measures showed promise for such applications by testing within-person change in biological age estimates calculated from biological samples taken when Study members were aged 26 years and again when they were aged 38 years. We computed change scores (age-38 value – age-26 value) to test how much telomere erosion actually took place over these 12 years and how many “ticks” were registered by the epigenetic clocks. (We did not test change in the KDM Biological Age and Age-related Homeostatic Dysregulation measures because the necessary data were not available at the age-26 assessment.)

Study members experienced an average of 0.15 (SD=0.30) T/S ratio units of telomere erosion over the 12-year follow-up. This telomere erosion was equivalent to about one-half of one standard deviation of the variance in telomere length at age 38 years. Study members’ epigenetic clocks ticked forward by 12-14 years (for the 353 CpG clock, M=12y, SD=3; for the 99 CpG clock, M=13y, SD=4; for the 71 CpG clock, M=14y, SD=5). This epigenetic “ticking” was equivalent to between 2 and 3 standard deviations of the variance in epigenetic clock values at age 38 years. We analyzed change in biological age estimated by Pace of Aging for comparison purposes. Because Pace of Aging estimates physiological-change-per-chronological-year, we multiplied each Study member’s Pace of aging by 12 to estimate change in biological age between chronological ages 26 and 38 years (M=12y, SD=5). Different measures showed similar distributions of variation (Figure 4).

**Figure 4.**
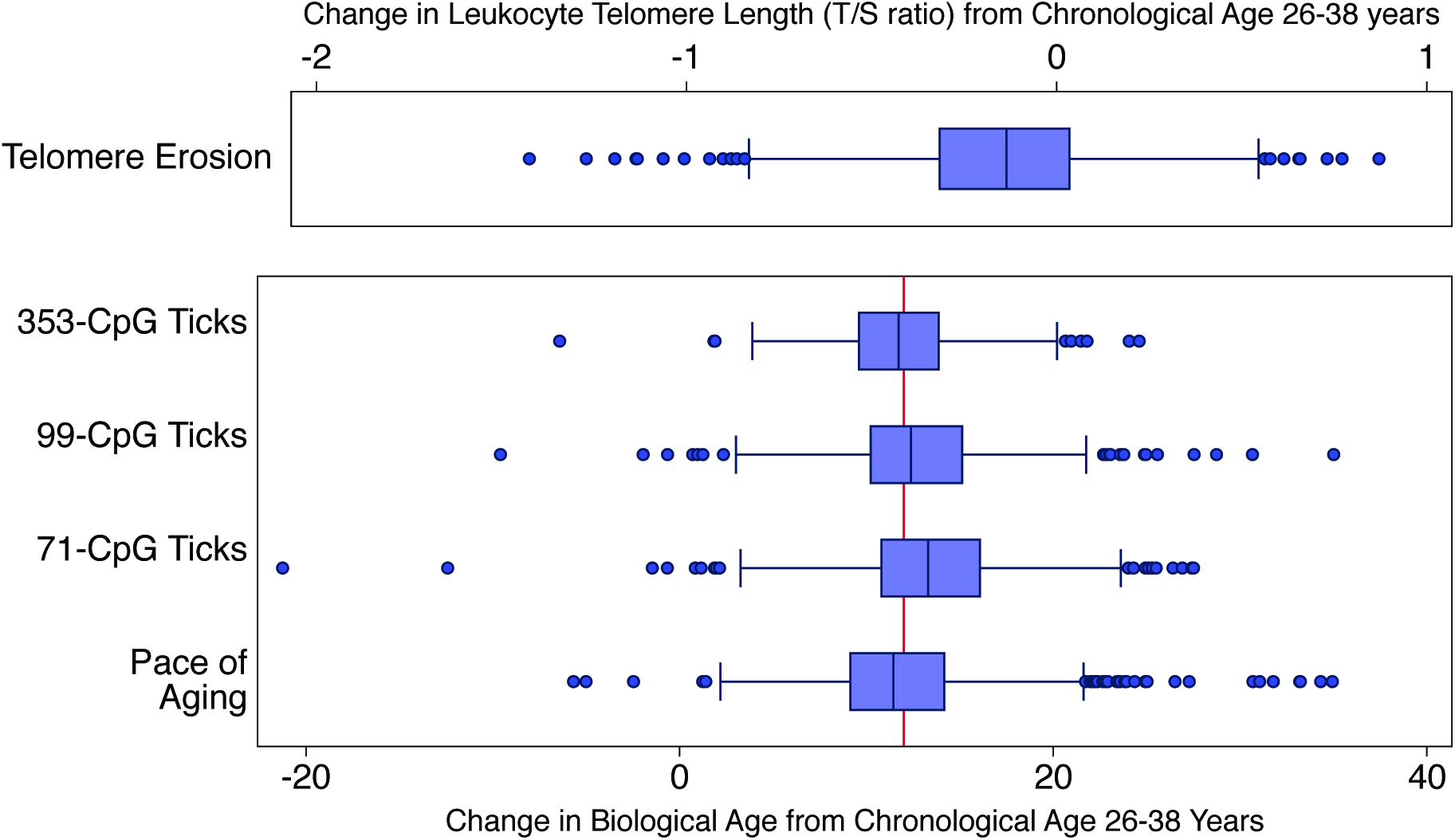
Changes in cross-sectional measures of biological aging between chronological ages 26 and 38 years in the Dunedin cohort. Telomere and epigenetic clock measurements were made from DNA samples extracted from peripheral blood collected when Study members were aged 26 and 38 years. Repeated observations of each individual were assayed together on the same plate/ methylation array to reduce batch effects. Telomere erosion and epigenetic ticking were measured by subtracting age-26 values from age-38 values. For comparison purposes, Pace of Aging is plotted alongside the epigenetic clocks. Pace of Aging is estimated from three repeated measurements at ages 26, 32, and 38 years of 18 different biomarkers. Pace of Aging is scaled in years of physiological change per chronological year. For this graph, Pace of Aging was multiplied by 12 to reflect the years of biological aging estimated to have occurred between ages 26 and 38 years. The vertical red line in the bottom panel of the figure indicates a value of 12 years, the actual amount of chronological time elapsed during the measurement interval.

To test if a common aging process influenced changes in different measures of biological aging, we computed correlations among change scores. Correlations among change scores showed a pattern similar to correlations among cross-sectional measures (Figure 5). Telomere erosion was not correlated with epigenetic ticking. Epigenetic ticking was correlated across the three different clocks (r=0.17-0.42). Epigenetic ticking was weakly correlated with Pace of Aging (r=0.06-0.09). The correlation between telomere erosion and Pace of aging was relatively high (r=0.24) because telomere erosion is a component of the Pace of Aging. When telomere erosion was excluded from Pace of Aging the correlation was reduced to near zero. Results were similar when Spearman correlations were computed to reduce the influence of extreme values (Supplemental Table 2).

**Figure 5.**
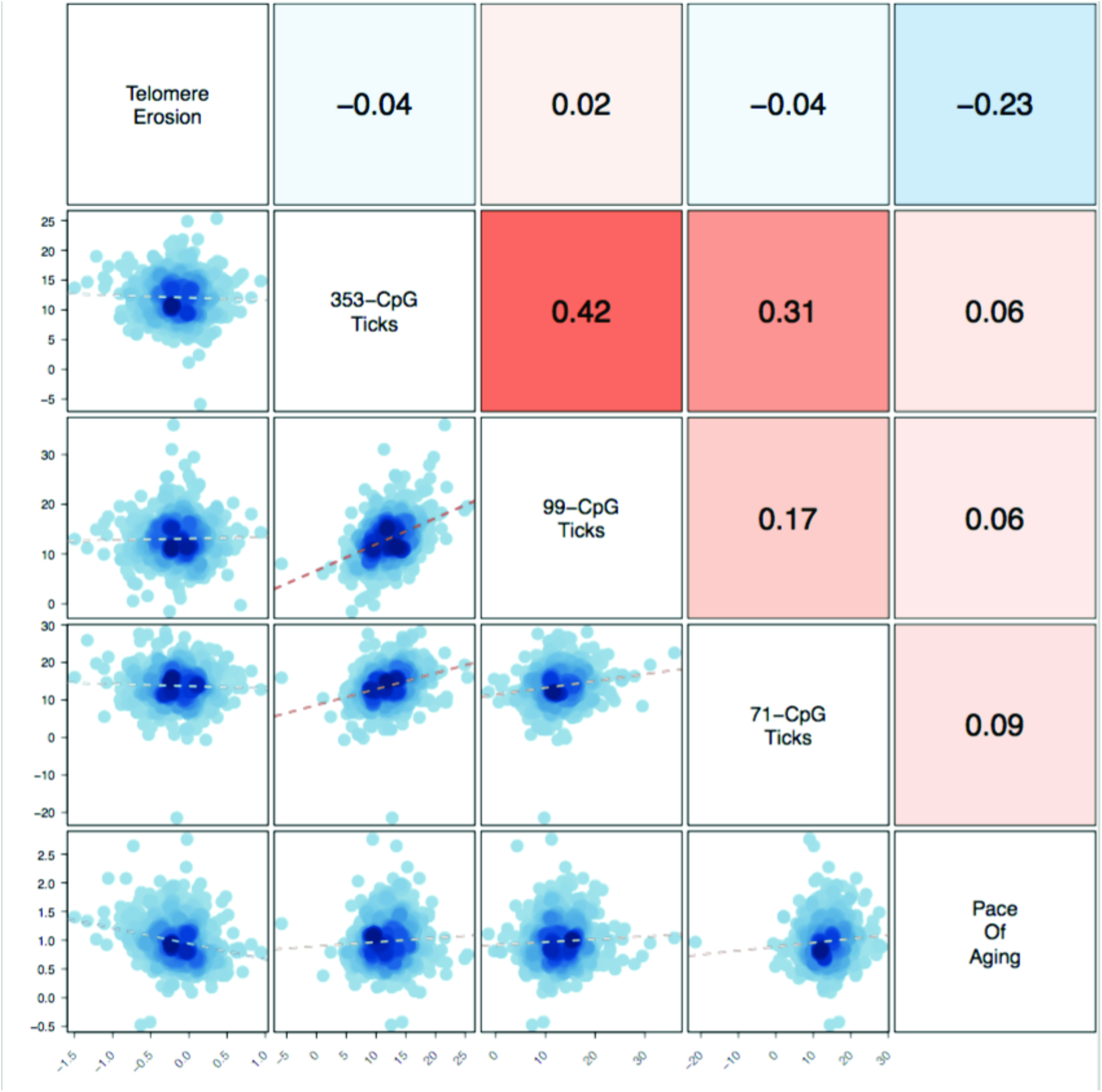
Relationships among longitudinal measures of biological aging. The figure shows a matrix of scatterplots and correlations illustrating relationships among 5 longitudinal measures of biological aging: telomere erosion, ticking of the 353-, 99-, and 71-CpG epigenetic clocks, and the Pace of Aging. Data are for n=733 Study members with complete data on all measures. Correlations are shown above the diagonal. (Correlations ≥0.07 are statistically significant at p<0.05.) Scatter plots are shown below the diagonal. Y-axis scales correspond to the biological aging metric listed to the left of the plot. X-axis scales correspond to the biological aging metric listed above the plot.

Change scores computed from repeated cross-sectional biological aging measures were not consistently associated with healthspan-related characteristics. Telomere erosion was not associated with healthspan-related characteristics (|r|≤0.04). Epigenetic ticking was also not associated with healthspan characteristics, with the exception of age-38 IQ score (r=0.11, p=0.003 for 353-CpG clock; r=0.09, p=0.017 for the 71-CpG clock) and self-rated health (r=-0.07, p=0.044 for the 71-CpG clock). Effect sizes are graphed in Figure 6 and Supplemental Figure 2.

**Figure 6.**
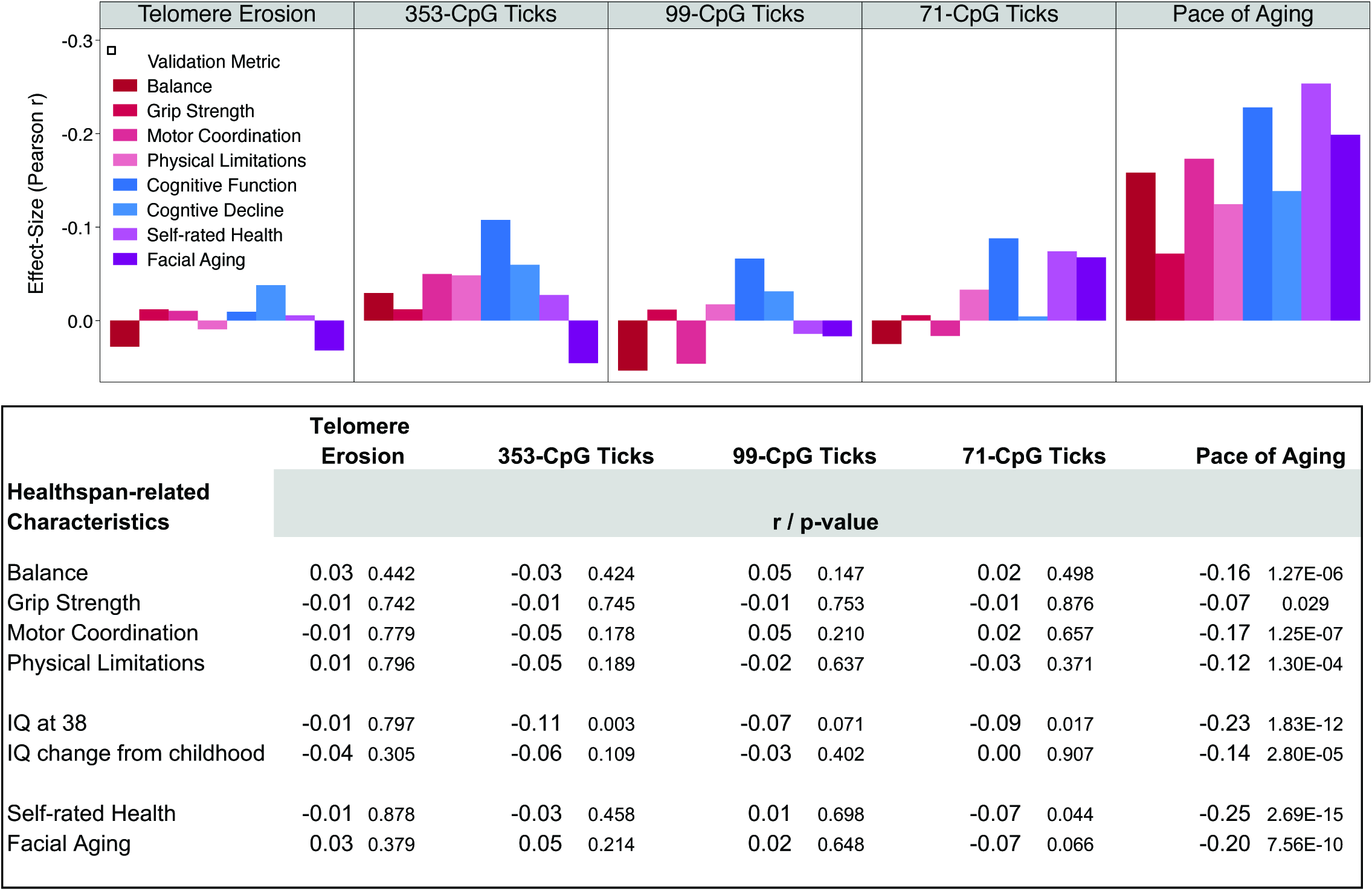
Associations of changes in cross-sectional biological aging measures and Pace of Aging with healthspan-related characteristics. The figure shows bar charts of effect-sizes for telomere erosion, ticking of 353-, 99-, and 71-CpG epigenetic clocks, and Pace of Aging. Effect-sizes were estimated for four measures of physical functioning (balance, grip strength, motor coordination, and self-reported physical limitations), cognitive functioning (IQ score at age 38 from the Wechsler Adult Intelligence Scale), cognitive decline (change in Wechsler-scale IQ score since childhood), and two measures of subjective aging (self-rated health and facial aging from assessments of facial photographs of the study member by independent raters). Validation metrics were scored so that higher values indicated increased healthspan. Telomere erosion was scored for this analysis so that higher values corresponded to more telomere erosion. Thus, the expected direction of association for all correlations was negative—because faster biological aging is expected to shorten healthspan. Standardized regression coefficients (interpretable as Pearson r) and their p-values are reported in the table below the figure.

A question about biological aging measures implemented during the middle period of the life course is whether they measure processes independent of weight gain^19,20^. To address this question, we repeated all tests of association between measures of biological aging and healthspan-related characteristics with statistical adjustment for body-mass index. Effect-sizes were essentially unchanged (Supplemental Tables 3-4).

## DISCUSSION

We studied seven proposed methods to quantify biological aging in a cohort of 964 individuals followed to midlife as part of the Dunedin Study. We quantified telomere length and erosion; 353-, 99-, and 71-CpG epigenetic clocks and their ticking rates; and three multi-biomarker algorithms: KDM Biological Age, Age-related Homeostatic Dysregulation, and the Pace of Aging. All of these measures indicated members of the Dunedin Study, despite being all the same chronological age, varied in their biological aging. Estimates of biological aging were in line with reports about these measures, e.g. epigenetic clocks varied around a mean of 38 years, matching the chronological age at which blood samples were taken. Moreover, when we compared Study members’ telomere and epigenetic clock measurements taken when they were aged 38 years with measurements from samples collected 12 years earlier, when they were aged 26 years, we detected the expected patterns of telomere erosion and epigenetic ticking. In fact, all three epigenetic clocks ticked forward by about 12 years, matching the amount of chronological time elapsed between sample collections. However, variation in different biological aging estimates did not appear to reflect a single aging process. Although epigenetic clocks correlated with one another and so did biomarker algorithms, correlations between the epigenetic clocks and biomarker algorithms were low, as were correlations of both sets of measures with telomere length. None of the measures of biological aging were strongly correlated with healthspan-related metrics of physical functioning, cognitive decline, or subjective aging. Telomere length and erosion and the 353- and 99-CpG epigenetic clocks and their ticking rates were not statistically associated with most healthspan-related metrics. The 71-CpG epigenetic clock generally showed statistically significant associations with healthspan-related metrics, although its ticking rate did not. Variability in results for the different epigenetic clocks may reflect differences in methods used to derive them^21,22^. KDM Biological Age, Age-related Homeostatic Dysregulation, and Pace of Aging generally showed statistically significant associations with healthspan-related metrics, although with small effect sizes.

This study had limitations. First, we studied a single birth cohort from New Zealand that lacked ethnic minority representation. Replication of findings in non-European cohorts is needed. Second, our follow-up extended only through age 38 years, precluding analysis of age-related disease, disability, and mortality. However, to evaluate geroprotective therapies that aim to prevent onset of age-related disease, biological aging measures that can quantify variation in comparatively young samples are needed. We tested associations of biological aging measures with measures of physical functioning, cognitive decline, and subjective signs of aging in our midlife cohort because these measures are prospectively related to disability and death and because they represent capacties geroprotective therapies will aim to preserve. Third, telomere erosion and epigenetic ticking measures were implemented using only two repeated measurements. Erosion and ticking measures thus could not separate measurement error from true change, as was possible with analysis of three repeated measures in Pace of Aging analysis. Studies of telomere erosion and epigenetic ticking over 3 or more repeated measurements are needed. Fourth, all molecular assays used to compute biological aging measures were implemented in samples from peripheral blood. Epigenetic clocks and telomeres may have different properties in other tissues^23^. However, as blood is among the most available tissues, biological aging measures that can be implemented in blood samples may be most suitable for translation to clinical trials of geroprotectors. Finally, our sample lacked power to detect very small effect-sizes. However, analyses were well-powered (>80%) to detect effect-sizes of r=0.1 and larger.

There is growing interest in methods to quantify processes of biological aging. These methods are needed for two purposes. One purpose is to serve as surrogate endpoints of healthspan extension in clinical trials of geroprotective therapies. Geroprotective therapies aim to slow the aging process and extend years of healthy life^24^. When clinical trials of such therapies are launched, the question remains: what should these trials study as outcomes? To measure healthspan and longevity will require follow-up intervals of several decades. Measures of biological aging that track the aging rate during and after administration of geroprotective therapy could allow for proof-of-concept human trials over much shorter follow-up intervals. A second purpose is to advance understanding of the biology of aging during the middle period of the life course, an understudied topic. Age-related diseases, physical frailty, and death are rare during this period. Biological aging can be quantified for everyone, substantially increasing power of studies hunting for genes, molecular processes, or psychosocial factors that influence fast, slow, or resilient aging during midlife.

Within this context, our study highlights progress, but also the need for a more systematic approach to development and testing of biological aging measures. Our findings do not imply that any particular measure of biological aging is better than the others nor that some or all of them are entirely unhelpful. For example, although we found no relationship between telomere length or epigenetic age and healthspan-related characteristics, there is evidence that these measures are associated with risk of disease and death in later life^25–28^. Conversely, although faster Pace of Aging predicted worse outcomes on the healthspan-related characteristics studied, its relation to mortality remains untested. To advance the geroscience agenda, biological aging research needs to address several gaps in knowledge. There are five main issues brought forward by our findings.

One issue is the chronological age of participants in biological aging studies. Indices of frailty already exist to quantify differences in older adults^29,30^. The greatest potential value of biological aging measures is in quantifying differences in humans who do not yet have age-related disease, most of whom are still middle aged or younger. Middle age is also when geroprotective therapies could in theory be most effective. So far, most studies have focused on older adults. Studies are needed of younger and mixed-age cohorts to compare performance of biological aging measures across periods of the lifespan, perhaps even including childhood^31^.

A second issue is the need for studies that compare different approaches to quantifying biological aging. Several methods to quantify biological aging have been put forward and have shown promise. Most studies so far concentrate on a single measure of biological aging or a single type of measure, e.g. studies have measured multiple epigenetic clocks^32,33^. Studies are needed that implement multiple methods in the same groups of humans in order to evaluate convergent and discriminant validity.

A third issue is the approach to validating biological age measures. The goal of geroscience is to extend healthspan. But validation studies of biological aging measures have focused primarily on predicting lifespan. Greater attention is needed to prediction of differences in the functional capacities that geroprotective therapies aim to preserve.

A fourth issue is how biological aging measures are developed in the first place. Chronological age is often used as the criterion standard for a biological aging measure^34^. But chronological age studied in cross-sectional data does not distinguish biological processes of aging from what are called “cohort effects”; older individuals were born and raised under different historical circumstances from younger ones^35^. Thus, chronological age may not provide an ideal criterion standard for biological aging. A related concern is mortality selection, the fact that comparatively fewer individuals from the earlier birth cohorts remain alive to be sampled at a given point in time^36^. Consequently, cross-sectional analyses of mixed-age samples may not be optimal for development of biological aging measures. Instead, longitudinal studies of within-individual change across repeated measures may provide a better platform for identification of biological changes specifically related to the aging process.

Finally, our findings highlight potentially important differences between biological aging measures implemented at different “levels” of analysis, as illustrated in Table 1. Telomere-length and epigenetic-clock methods are cellular-level measures implemented in our study in only a single tissue, blood. In contrast, the KDM Biological Age, Age-related Homeostatic Dysregulation, and Pace of Aging measures draw information from multiple systems throughout the body. It is possible that composite measures of, e.g. epigenetic-clocks, from multiple tissues might show stronger correlation with the other measures of aging and with the healthspan-related characteristics we studied. Quantifications of biological aging that can be implemented at the level of a single cell are appealing because they allow for direct investigation of cellular-level mechanisms of aging. However, for the purposes of measuring effectiveness of geroprotective therapies, quantifications of biological aging that draw information from multiple bodily systems may be more sensitive and specific with respect to the target outcome of healthspan extension. Based on our analysis, it is possible that a geroprotective therapy might retard one measure of aging, but fail to produce any healthspan extension as ascertained by other measures, leaving efficacy of the therapy in question.

Methods to quantify biological aging have potential to advance efforts to elucidate the basic biology of aging and to translate emerging geroprotective therapies from animals to humans. Quantifications of biological aging may also provide clinicians with a tool to communicate complex health information to patients in an easy-to-understand way. Finally, biological age measures can provide a tool for precision medicine, helping physicians decide when a patient should begin screening for age-related conditions. To realize this promise, efforts are needed to harmonize research practices for testing proposed biological aging measures. Research on biological aging recently experienced a growth spurt. As new measures are subjected to increasingly stringent tests, discoveries will be tempered by caveats. Rather than discouraging further investigation, these caveats should be interpreted as signs of maturation and encourage redoubled efforts to develop measures of biological aging.

## METHODS

### Sample

Participants are members of the Dunedin Study, a longitudinal investigation of health and behavior in a complete birth cohort. Study members (N=1,037; 91% of eligible births; 52% male) were all individuals born between April 1972 and March 1973 in Dunedin, New Zealand (NZ), who were eligible based on residence in the province and who participated in the first assessment at age 3. The cohort represented the full range of socioeconomic status in the general population of New Zealand’s South Island. On adult health, the cohort matches the NZ National Health & Nutrition Survey (e.g., BMI, smoking, GP visits)^37^. Cohort members are primarily white; fewer than 7% self-identify as having partial non-Caucasian ancestry, matching the South Island^37^. Assessments were carried out at birth and ages 3, 5, 7, 9, 11, 13, 15, 18, 21, 26, 32, and, most recently, 38 years, when 95% of the 1,007 study members still alive took part. At each assessment, each study member is brought to the research unit for a full day of interviews and examinations. The Otago Ethics Committee approved each phase of the study and informed consent was obtained from all study members.

### Quantification of Biological Aging

We implemented cross-sectional biological age measures using data collected when Dunedin Study members were aged 38 years. We constructed longitudinal biological aging measures from repeated cross-sectional assessments of telomere length and epigenetic clocks when Study members were aged 26 and 38 years. We measured the longitudinal Pace of Aging from repeated biomarker assessments at ages 26, 32, and 38 years. Data to quantify at least one biological aging measure were available on N=964 individuals. Measures are described in detail below.

#### Telomere length

Telomere length was measured from leukocyte DNA collected at ages 26 and 38 years. Leukocyte DNA was extracted from blood using standard procedures^38,39^. DNA was stored at -80°C. All DNA samples were assayed for leukocyte telomere length at the same time. Leukocyte telomere length was measured using a validated quantitative PCR method^40^, as previously described^41^, which determines mean telomere length across all chromosomes for all cells sampled. The method involves two quantitative PCR reactions for each subject; one for a single-copy gene (S) and the other in the telomeric repeat region (T). All DNA samples were run in triplicate for telomere and single-copy reactions.

Measurement artifacts (e.g., differences in plate conditions) may lead to spurious results when comparing leukocyte telomere length measured on the same individual at different ages. To eliminate such artifacts, we assayed DNA triplicates from the same individual from all time points, on the same plate. CV for triplicate Ct values was 0.81% for the telomere (T) and 0.48% for the single-copy gene (S). We computed change in telomere length as the Age-38 T/S ratio – Age-26 T/S ratio. Telomere data were available for N=829 Study members at age 38, for N=812 Study members at age 26, and for N=758 Study members at both ages of measurement.

#### Epigenetic Clocks

Epigenetic clocks were calculated using leukocyte DNA collected at ages 26 and 38 years. 500ng of DNA from each sample was treated with sodium bisulfite, using the EZ-96 DNA Methylation kit (Zymo Research, CA, USA). DNA methylation was quantified using the Illumina Infinium HumanMethylation450 BeadChip (Illumina Inc, CA, USA) run on an Illumina iScan System (Illumina, CA, USA) using the manufacturers’ standard protocol. Briefly, these arrays simultaneously interrogate >485,000 methylation sites distributed across the genome. Samples were arranged into 96-well plates so that within-individual age-26 and -38 DNA samples were hybridized in the same row of the arrays (i.e. age 26 and 38 DNA samples from the same individual occupy array columns 1 and 2 of the same row). Array analysis was performed by the Duke University Molecular Physiology Institute Genomics Core Facility using the iScan platform (Illumina). Data quality control and normalization was carried out using the *Methylumi* Bioconductor package in the R statistical programming environment.

We analyzed three epigenetic clocks. The first clock, proposed by Horvath, included 353 CpG sites^9^. The second clock, proposed by Hannum and colleagues, included 71 CpG sites^10^. The third clock, proposed by Weidner and colleagues, included 99 CpG sites^13,42^. Study members’ epigenetic clock values for the 353-CpG and 71-CpG clocks were calculated using Horvath’s website (https://labs.genetics.ucla.edu/horvath/dnamage/). Epigenetic clock values for the 99-CpG clock were calculated using the algorithm published by the Wagner lab^43,32^. Epigenetic clock values were available for N=818 Study members at age 38, for N=821 Study members at age 26, and for N=743 Study members at both ages of measurement.

#### Biological Age

As described previously^14^, we calculated each Study member’s Biological Age at age 38 years using the Klemera-Doubal equation^34^ and parameters estimated from the NHANES-III dataset^16^ for ten biomarkers: Glycated hemoglobin, Forced expiratory volume in one second (FEV1), Blood pressure (systolic), Total cholesterol, C-reactive protein, Creatinine, Urea nitrogen, Albumin, Alkaline phosphatase, and Cytomegalovirus IgG. Data to calculate Biological Age data were available for N=904 Study members.

#### Age-Related Homeostatic Dysregulation

We measured age-related homeostatic dysregulation by applying the biomarker Mahalanobis distance method described by Cohen and colleagues^12,17,44^ to Study members’ age-38 biomarker values. The biomarker Mahalanobis distance method measures how aberrant an individual’s physiology is relative to a reference norm^12^. Cohen and colleagues used chronologically young individuals to form this reference norm for their calculations^17^. They interpreted biomarker Mahalanobis distance from the reference as an indicator of age-related homeostatic dysregulation, a sign of biological aging. We formed our reference from the Dunedin Study members’ biomarker values at age 26 years, the youngest age at which the biomarkers were measured. Thus, a Study member’s biomarker Mahalanobis distance quantifies homeostatic dysregulation relative to the cohort’s age-26 norm. We calculated Mahalanobis distance based on 18 biomarkers with repeated measures at ages 26 and 38 years (the same 18 biomarkers we previously used to compute Study members’ Pace of Aging^14^, see below). Distances were log transformed for analysis. Age-related Homeostatic Dysregulation was measured for N=954 Study members.

#### Pace of Aging

As described previously^14^, we measured Pace of Aging with repeated assessments of a panel of 18 biomarkers taken at ages 26, 32, and 38 years. The biomarkers were: Apolipoprotein B100/A1 ratio, Blood pressure (mean arterial pressure), Body mass index (BMI) and Waist-hip ratio, C-reactive protein and white blood cell count, Cardiorespiratory fitness (VO_2_Max), Creatinine clearance, Forced expiratory volume in one second (FEV_1_) and Forced vital capacity ratio (FEV_1_/FVC), Glycated hemoglobin, High density lipoprotein (HDL), Lipoprotein(a), Leukocyte telomere length (LTL), Periodontal disease, Total cholesterol, Triglycerides, Urea nitrogen. For each biomarker, we calculated the Study member’s personal rate of change using mixed-effects growth models. We combined these rates of change into a single index scaled in years of physiological change occurring per one chronological year. The average Study member had Pace of Aging equal to one year of physiological change per one chronological year. The fastest-aging Study members experienced more than twice that rate of physiological change. The slowest-aging study members experienced almost no change at all. Pace of Aging was measured for N=954 Study members.

### Healthspan-related Characteristics

Healthspan-related characteristics were measured when Study members were aged 38 years. All healthspan-related characteristics were transformed to sex-specific Z-scores for analysis, with the exception of cognitive test scores and the facial aging measure, which are sex neutral.

#### Physical Functioning

We employed three measures. First, we measured balance as the maximum time achieved across three trials of the Unipedal Stance Test (with eyes closed)^45–47^. Second, we measured grip strength with dominant hand (elbow held at 90°, upper arm held tight against the trunk) as the maximum value achieved across three trials using a Jamar digital dynamometer^48,49^. Third, we measured motor functioning as the time to completion of the Grooved Pegboard Test with the dominant hand^50^.

#### Physical Limitations

Study member responses (“limited a lot,” “limited a little,” “not limited at all”) to the 10-item SF-36 physical functioning scale^51^ assessed their difficulty with completing various activities, e.g., climbing several flights of stairs, walking more than 1 km, participating in strenuous sports.

#### Cognitive Testing

IQ is a highly reliable measure of general intellectual functioning that captures overall ability across differentiable cognitive functions. We measured IQ from the individually administered Wechsler Intelligence Scale for Children-Revised (WISC-R; averaged across ages 7, 9, 11, and 13)^52^ and the Wechsler Adult Intelligence Scale-IV (WAIS-IV; age 38)^53^, both with M=100 and SD=15. We measured IQ decline by comparing scores from the WISC-R (in childhood) and the WAIS-IV (at age 38 years). Analyses of subtests are reported in the **Supplementary Information**.

#### Self-Rated Health

Study members rated their health on a scale of 1-5 (poor, fair, good, very good, or excellent).

#### Facial aging

We took two measurements of perceived age based on facial photographs. First, Age Range was assessed by an independent panel of 4 Duke University undergraduate raters. Raters were presented with standardized (non-smiling) facial photographs of Study members (taken with a Canon PowerShot G11 camera with an optical zoom, Canon Inc., Tokyo, Japan) and were kept blind to their actual age. Photos were divided into sex-segregated slideshow batches containing approximately 50 photos, viewed for 10s each. Raters were randomized to viewing the slideshow batches in either forward progression or backwards progression. They used a Likert scale to categorize each Study member into a 5-year age range (i.e., from 20-24 years old up to 65-70 years). Scores for each study member were averaged across all raters (α=0.71). The second measure, Relative Age, was assessed by a different panel of 4 Duke University undergraduate raters. The raters were told that all photos were of people aged 38 years old. Raters then used a 7-item Likert scale to assign a “relative age” to each study member (1=“young looking”, 7=“old looking”). Scores for each study member were averaged across all raters (α=0.72). Age Range and Relative Age were highly correlated (r=0.73). To derive a measure of perceived age at 38 years, we standardized and averaged both Age Range and Relative Age scores to create Facial Age at 38 years.

## ACKNOWLEDGEMENT

We thank the Dunedin Study members, their parents, teachers, partners, and peer informants, and Study founder Phil Silva. The Dunedin Multidisciplinary Health and Development Research Unit is supported by the New Zealand Health Research Council and New Zealand Ministry of Business, Innovation and Employment (MBIE). This research received support from US-National Institute of Aging grants R01AG032282, R01AG048895, and 1R01AG049789, UK Medical Research Council grant MR/K00381X, and UK ESRC grant ES/M010309/1. Additional support was provided by P30AG028716, P30AG034424, and by the Jacobs Foundation. DWB is supported by an Early-Career Research Fellowship from the Jacobs Foundation. AAC is a member of the FRQS-funded Centre de recherche sur le vieillissement and Centre de recherche du CHUS, and holds a CIHR New Investigator Salary Award.

## SUPPLEMENTARY INFORMATION

Supplement to Telomere, epigenetic clock, and biomarker-composite quantifications of biological aging: Do they measure the same thing?

DW Belsky et al.

**Supplemental Table 1.** Relationships among seven measures of biological aging in a birth cohort at chronological age 38 years – Spearman correlations.

|  |  | <u>Spearman correlations</u> |  |  |  |  |  | <u>p-values for Spearman correlations</u> |  |  |  |  |  |
| --- | --- | --- | --- | --- | --- | --- | --- | --- | --- | --- | --- | --- | --- |
|  |  | (1) | (2) | (3) | (4) | (5) | (6) | (1) | (2) | (3) | (4) | (5) | (6) |
| **Spearman Correlations** |  |  |  |  |  |  |  |  |  |  |  |  |  |
| (1) | Telomere Length |  |  |  |  |  |  |  |  |  |  |  |  |
| (2) | 353-CpG Clock | -0.05 |  |  |  |  |  | 0.174 |  |  |  |  |  |
| (3) | 99-CpG Clock | -0.04 | 0.53 |  |  |  |  | 0.282 | 5.66E-60 |  |  |  |  |
| (4) | 71-CpG Clock | -0.04 | 0.41 | 0.34 |  |  |  | 0.276 | 3.06E-33 | 5.38E-23 |  |  |  |
| (5) | KDM Biological Age | -0.05 | 0.12 | 0.08 | 0.14 |  |  | 0.152 | 0.001 | 0.028 | 6.52E-05 |  |  |
| (6) | Age-related Homeostatic Dysregulation | 0.02 | 0.04 | 0.03 | 0.09 | 0.40 |  | 0.652 | 0.272 | 0.390 | 0.013 | 8.35E-32 |  |
| (7) | Pace of Aging | -0.03 | 0.00 | -0.01 | 0.12 | 0.36 | 0.48 | 0.351 | 0.905 | 0.711 | 0.001 | 1.48E-25 | 5.20E-47 |

**Supplemental Table 2.**
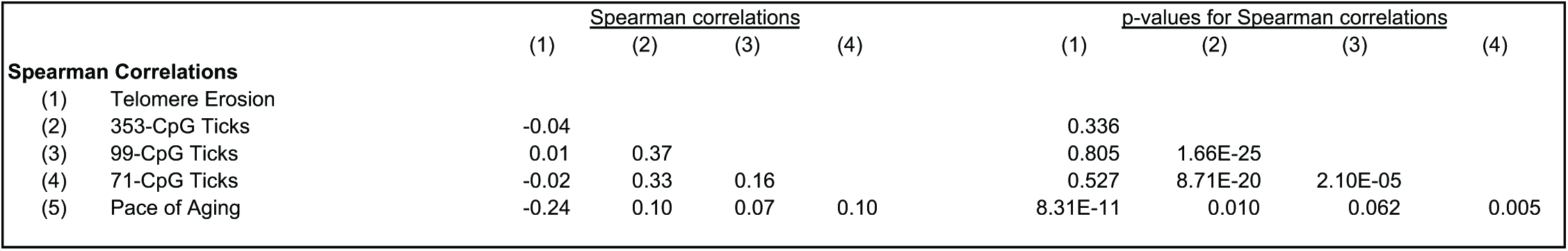
Relationships among longitudinal measures of biological aging – Spearman correlations.

|  |  | <u>Spearman correlations</u> |  |  |  | <u>p-values for Spearman correlations</u> |  |  |  |
| --- | --- | --- | --- | --- | --- | --- | --- | --- | --- |
|  |  | (1) | (2) | (3) | (4) | (1) | (2) | (3) | (4) |
| **Spearman Correlations** |  |  |  |  |  |  |  |  |  |
| (1) | Telomere Erosion |  |  |  |  |  |  |  |  |
| (2) | 353-CpG Ticks | -0.04 |  |  |  | 0.336 |  |  |  |
| (3) | 99-CpG Ticks | 0.01 | 0.37 |  |  | 0.805 | 1.66E-25 |  |  |
| (4) | 71-CpG Ticks | -0.02 | 0.33 | 0.16 |  | 0.527 | 8.71E-20 | 2.10E-05 |  |
| (5) | Pace of Aging | -0.24 | 0.10 | 0.07 | 0.10 | 8.31E-11 | 0.010 | 0.062 | 0.005 |

**Supplemental Figure 1.**
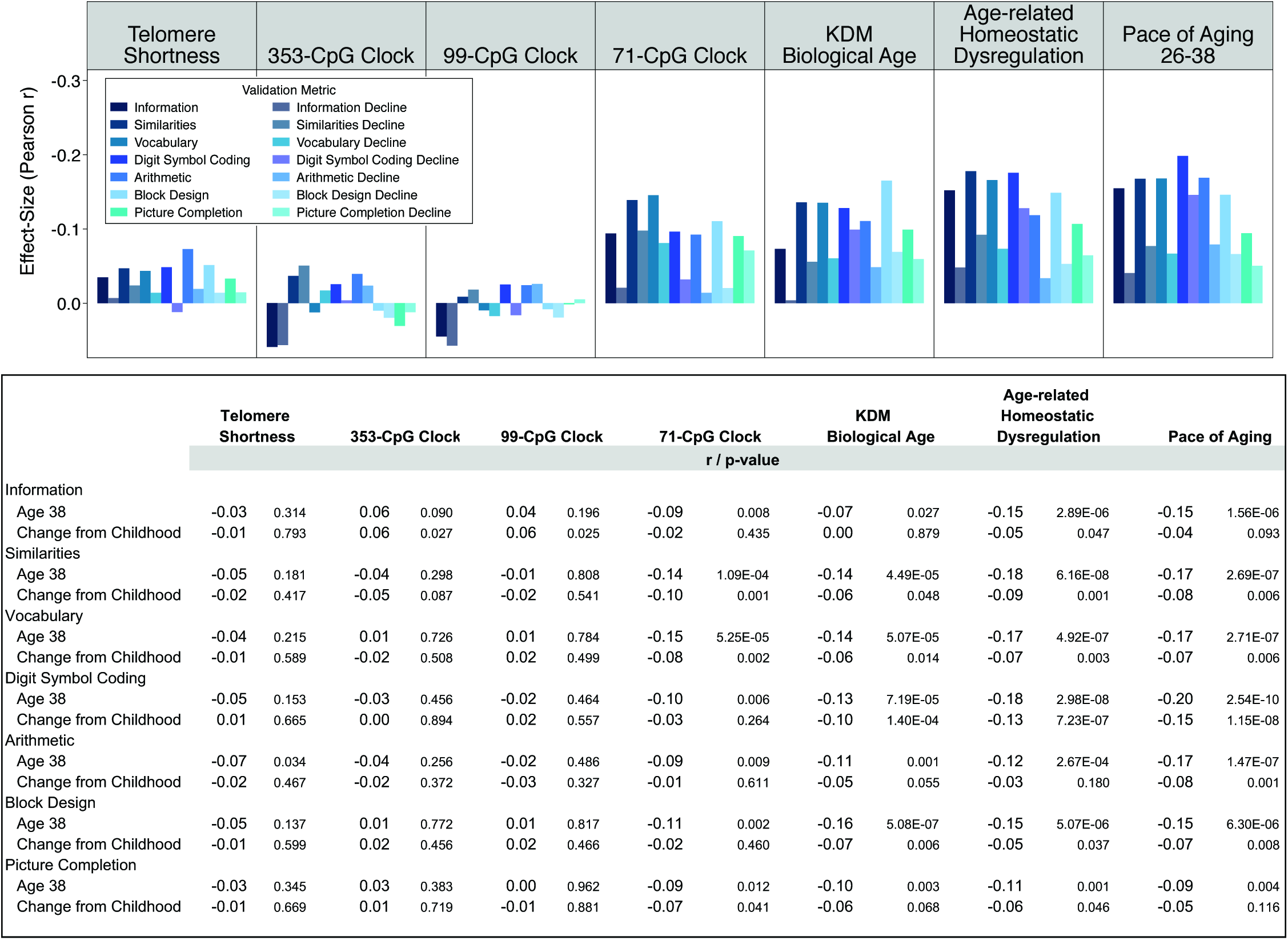
Associations of cross-sectional biological aging measures and Pace of Aging with subtests of cognitive functioning and cognitive decline. The figure shows bar charts of effect-sizes (Pearson r)for each of the seven measures of biological aging. Effect-sizes were estimated for seven tests of cognitive function administered in parallel during childhood and age-38 assessments. The tests were subscales of the Wechsler Intelligence Tests. There were three tests of so-called “crystalized” cognitive functions (Information, Similarities, and Vocabulary), and four tests of so-called “fluid” cognitive functions (Digit Symbol Coding, Arithmetic, Block Design, and Picture Completion). All tests were scored so that higher values corresponded to indication of better cognitive functioning. Telomere length was reversed for this analysis so that higher values corresponded to shorter telomeres. Thus, the expected direction of association for all correlations was negative—because faster biological aging is expected to hasten cognitive decline. Standardized regression coefficients (interpretable as Pearson r) and their p-values are reported in the table below the figure. For each test, the graph plots the effect-size for association between biological aging and age-38 test performance first (darker shaded bars), followed by the effect-size for association between biological aging and actual decline in test performance between childhood and age 38 (lighter shaded bars).

**Supplemental Figure 2.**
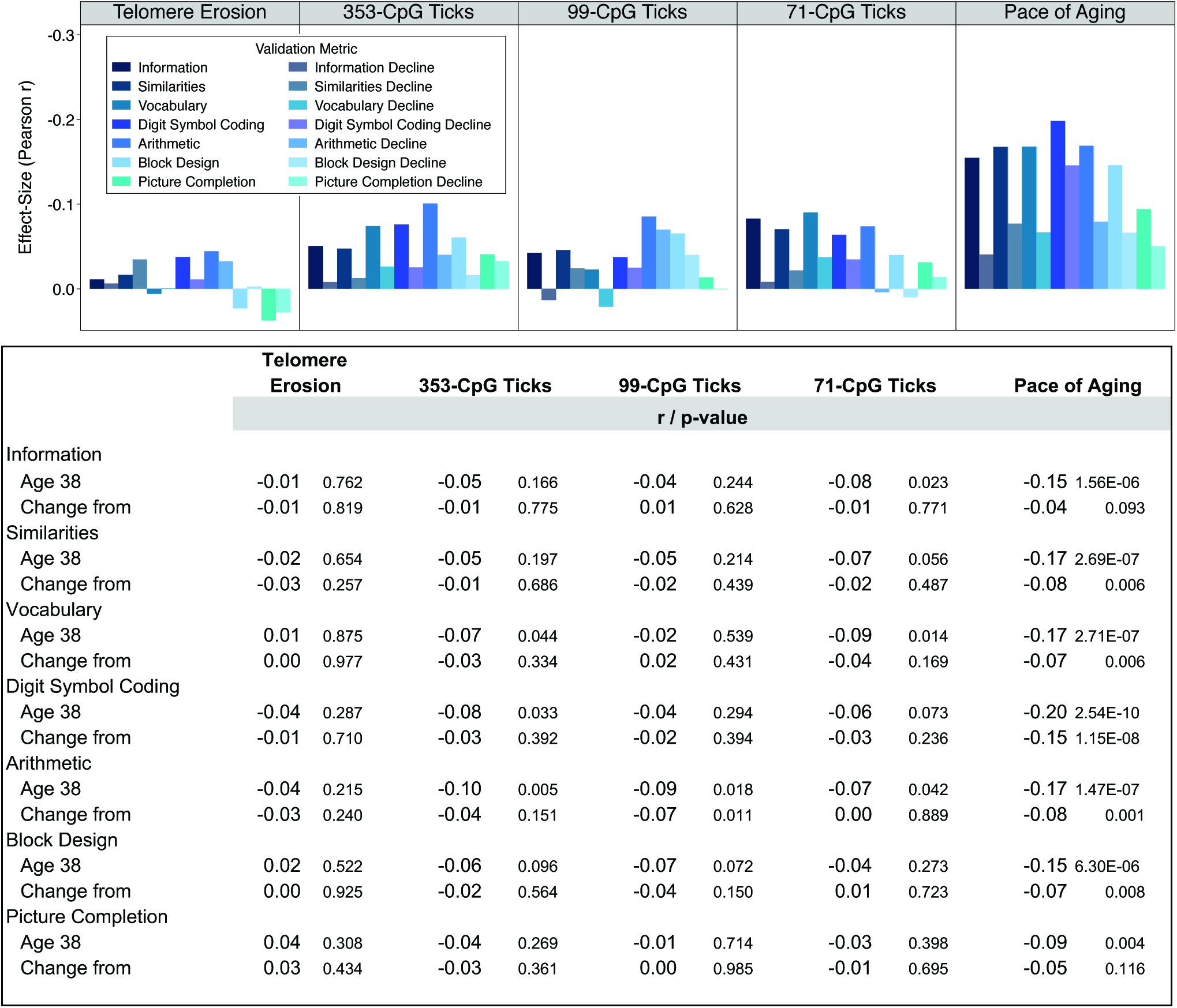
Associations of changes in cross-sectional biological aging measures and Pace of Aging with subtests of cognitive functioning and cognitive decline. The figure shows bar charts of effect-sizes (Pearson r) for telomere erosion, ticking of 353-, 99-, and 71-CpG epigenetic clocks, and Pace of Aging. Effectsizes were estimated for seven tests of cognitive function administered in parallel during childhood and age-38 assessments. The tests were subscales of the Wechsler Intelligence Tests. There were three tests of so-called “crystalized” cognitive functions (Information, Similarities, and Vocabulary), and four tests of so-called “fluid” cognitive functions (Digit Symbol Coding, Arithmetic, Block Design, and Picture Completion). All tests were scored so that higher values corresponded to indication of better cognitive functioning. Telomere erosion was scored for this analysis so that higher values corresponded to more telomere erosion. Thus, the expected direction of association for all correlations was negative—because faster aging is expected to hasten cognitive decline. Standardized regression coefficients (interpretable as Pearson r) and their p-values are reported in the table below the figure. For each test, the graph plots the effect-size for association between biological aging and age-38 test performance first (darker shaded bars), followed by the effect-size for association between biological aging and actual decline in test performance between childhood and age 38 (lighter shaded bars).

**Supplemental Table 3.**
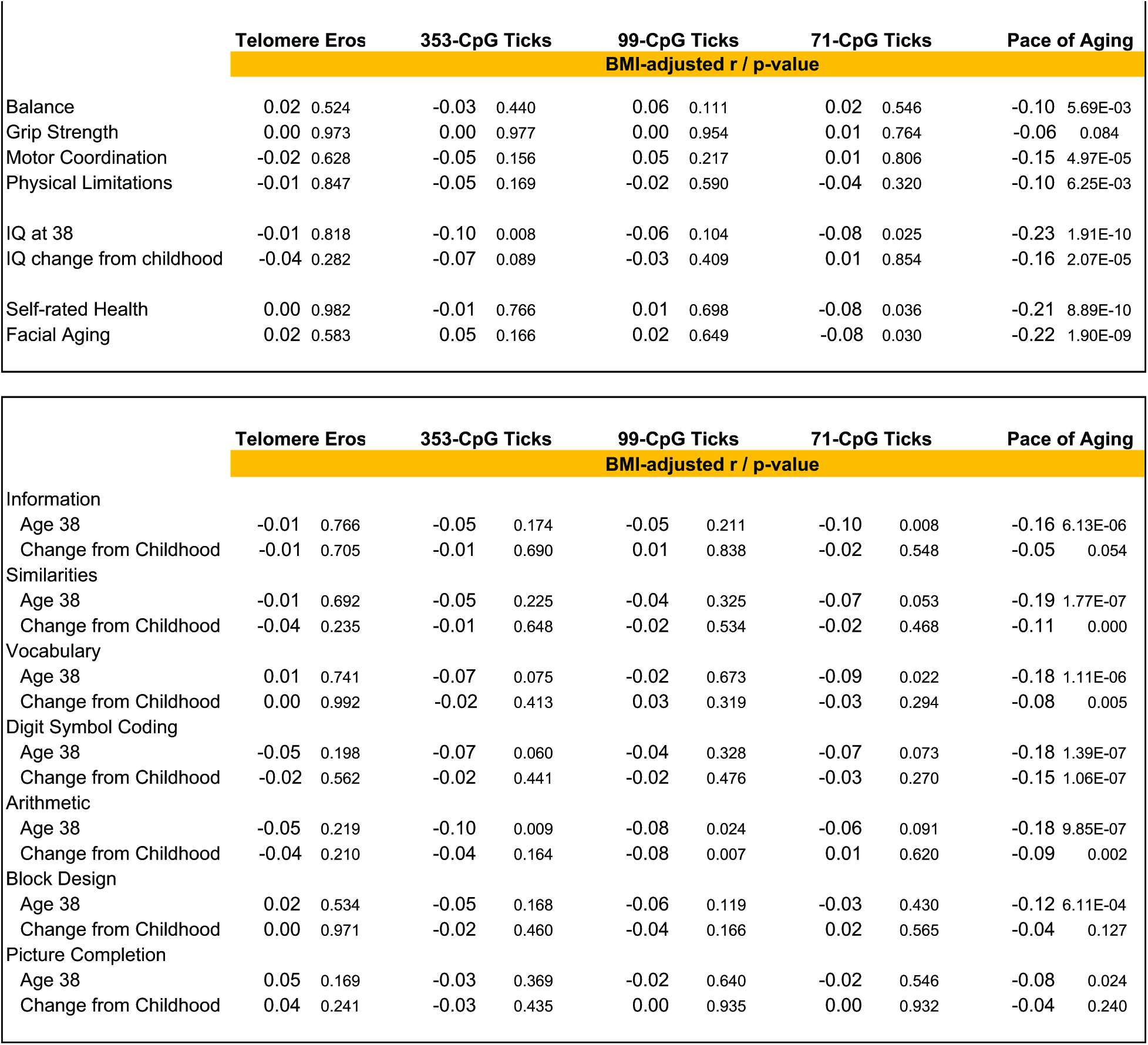
Associations of cross-sectional biological aging measures and Pace of Aging with healthspan-related characteristics & cognitive subtests after adjustment for bodymass index.

**Supplemental Table 4.**
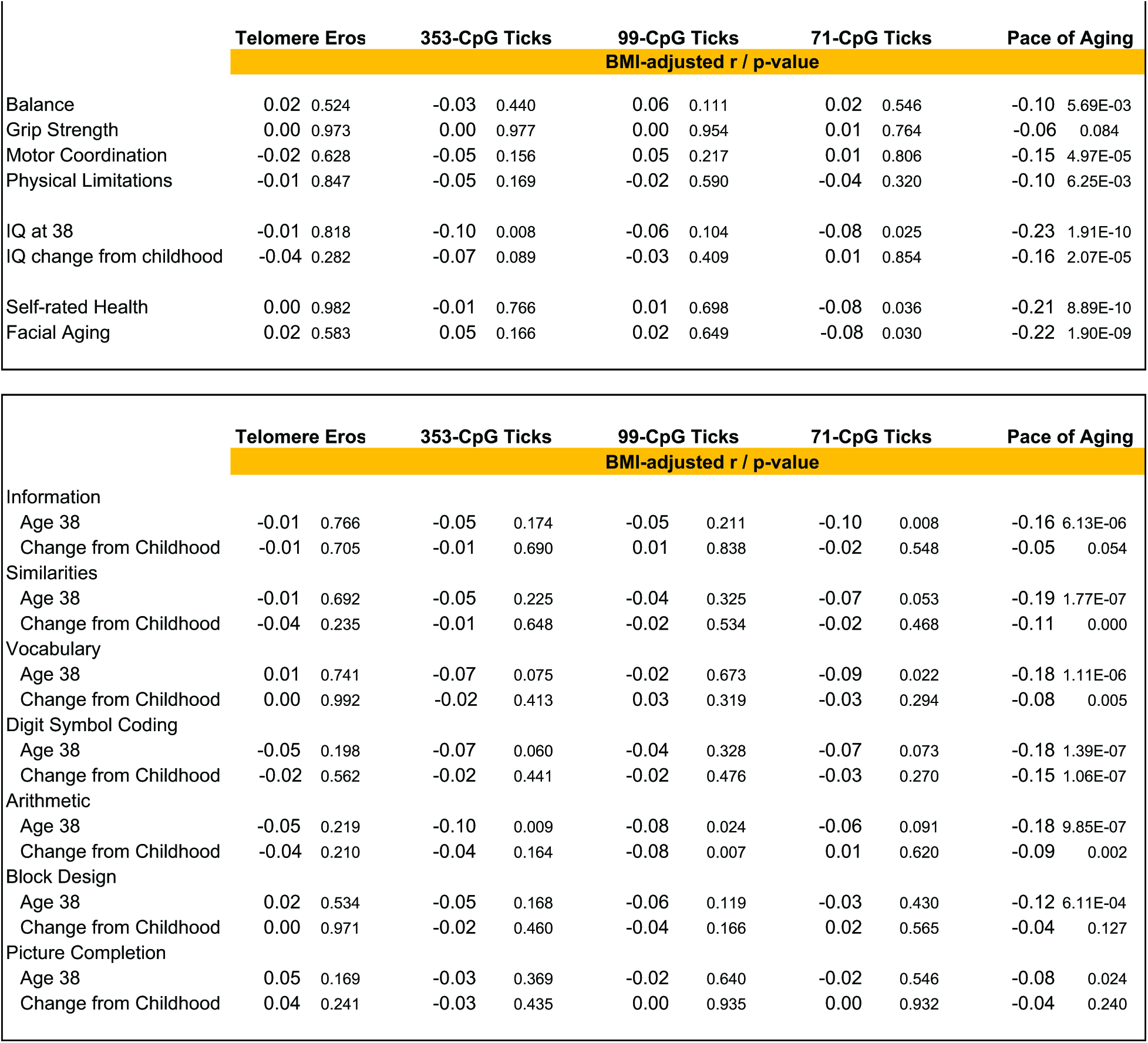
Associations of changes in cross-sectional biological aging measures and Pace of Aging with healthspan-related characteristics and cognitive subtests after adjustment for change in body mass index.

